# No consistent evidence for microbiota in murine placental and fetal tissues

**DOI:** 10.1101/2019.12.10.872275

**Authors:** Kevin R. Theis, Roberto Romero, Jonathan M. Greenberg, Andrew D. Winters, Valeria Garcia-Flores, Kenichiro Motomura, Madison M. Ahmad, Jose Galaz, Marcia Arenas-Hernandez, Nardhy Gomez-Lopez

**Affiliations:** Department of Biochemistry, Microbiology and Immunology, Wayne State University School of Medicine, Detroit, MI, USA; Perinatology Research Branch, Division of Obstetrics and Maternal-Fetal Medicine, Division of Intramural Research, *Eunice Kennedy Shriver* National Institute of Child Health and Human Development, National Institutes of Health, U.S. Department of Health and Human Services. Bethesda, MD and Detroit, MI; Department of Obstetrics and Gynecology, University of Michigan, Ann Arbor, MI, USA; Department of Epidemiology and Biostatistics, Michigan State University, East Lansing, MI, USA; Center for Molecular Medicine and Genetics, Wayne State University, Detroit, MI, USA; Detroit Medical Center, Detroit, MI, USA; Department of Obstetrics and Gynecology, Florida International University, Miami, FL, USA; Department of Obstetrics and Gynecology, Wayne State University School of Medicine, Detroit, MI, USA; Department of Obstetrics and Gynecology, Faculty of Medicine, Pontificia Universidad Católica de Chile, Santiago, Chile; Department of Physiology, Wayne State University School of Medicine, Detroit, Michigan, USA

**Keywords:** Microbiome, low microbial biomass sample, pregnancy, *in utero* colonization, mouse model

## Abstract

The existence of a placental microbiota and *in utero* colonization of the fetus has been the subject of recent debate. The objective of this study was to determine whether the placental and fetal tissues of mice harbor bacterial communities. Bacterial profiles of the placenta and fetal brain, lung, liver, and intestine were characterized through culture, qPCR, and 16S rRNA gene sequencing. These profiles were compared to those of the maternal mouth, lung, liver, uterus, cervix, vagina, and intestine, as well as to background technical controls. Positive bacterial cultures from placental and fetal tissues were rare; of the 165 total bacterial cultures of placental tissues from the 11 mice included in this study, only nine yielded at least a single colony, and five of those nine positive cultures came from a single mouse. Cultures of fetal intestinal tissues yielded just a single bacterial isolate: *Staphylococcus hominis*, a common skin bacterium. Bacterial loads of placental and fetal brain, lung, liver, and intestinal tissues were not higher than those of DNA contamination controls and did not yield substantive 16S rRNA gene sequencing libraries. From all placental or fetal tissues (N = 49), there was only a single bacterial isolate that came from a fetal brain sample having a bacterial load higher than that of contamination controls and that was identified in sequence-based surveys of at least one of its corresponding maternal samples. Therefore, using multiple modes of microbiologic inquiry, there was not consistent evidence of bacterial communities in the placental and fetal tissues of mice.

**IMPORTANCE:** The prevailing paradigm in obstetrics has been the sterile womb hypothesis, which posits that fetuses are first colonized by microorganisms during the delivery process. However, some are now suggesting that fetuses are consistently colonized by microorganisms *in utero* by microbial communities that inhabit the placenta and intra-amniotic environment. Given the established causal role of microbial invasion of the amniotic cavity (i.e. intra-amniotic infection) in pregnancy complications, especially preterm birth, if the *in utero* colonization hypothesis were true, there are several aspects of current understanding that will need to be reconsidered including the magnitude of intra-amniotic microbial load required to cause disease and their potential influence on the ontogeny of the immune system. However, acceptance of the *in utero* colonization hypothesis is premature. Herein, we do not find consistent evidence for placental and fetal microbiota in mice using culture, qPCR, and DNA sequencing.

## INTRODUCTION

The existence of resident microbial communities in the placenta (1-30) and potentially of *in utero* microbial colonization of the fetus (6, 22, 24, 31-33) have been the subject of recent debate. A few studies of the human placenta have reported the consistent detection of bacteria through microscopy (20, 34, 35) or culture (6). However, many recent studies detecting the presence of bacteria in the placenta, and thus proposing the existence of a placental microbiota, have done so using DNA sequencing techniques (1-5, 8-12, 15).

A principal caveat of these studies has been that the detected bacteria may reflect background DNA contamination from DNA extraction kits and PCR reagents rather than resident bacterial communities within the placenta (7, 14, 17, 21, 25). Another critique has been that even if the detected molecular signals of bacteria in the placenta are not background DNA contaminants, they may nevertheless reflect bacterial products circulating in the maternal blood rather than viable bacterial communities inhabiting the placenta (36). This is important because the detection of microbial DNA is not the same as the identification of a viable microorganism, and fetal exposure to microbial products (37), including DNA (38-40), is not commensurate with *in utero* microbial colonization of the fetus (24).

To establish the existence of resident bacterial communities in placental or fetal tissues would require: 1) the identification of bacterial DNA in placental or fetal tissues that is distinct from bacterial DNA detected in technical controls (e.g. DNA extraction kits, PCR reagents, laboratory environments), 2) confirmation that the bacterial load of placental or fetal tissues exceeds that of technical controls through quantitative real-time PCR (qPCR), 3) visualization of bacteria in placental or fetal tissues using microscopy, 4) demonstration of the viability of bacteria in these tissues through culture, and 5) ecological plausibility (i.e. the detected bacteria could survive and reproduce in these tissues) (21, 25). To date, these criteria have not been met in any one study, and affirmative conclusions about the existence of a placental microbiota and *in utero* microbial colonization of the fetus are premature.

While much of the debate regarding the existence of resident microbial communities in the placenta and of *in utero* microbial colonization of the fetus has focused on humans, a few studies have been conducted on mammalian animal models as well. Specifically, studies of the placental and fetal tissues of rats and mice (22, 33, 40), and preliminary studies of the placental and fetal tissues of rhesus macaques (41-45), have suggested that these tissues harbor bacterial communities. The benefit of using animal models to investigate the existence of *in utero* microbiota is that you can surgically obtain placental and fetal tissues prior to the onset of labor – if the fetal tissues are populated by viable bacterial communities, then bacterial colonization of the fetus had to occur *in utero*.

The objective of the current study was to determine whether the placental and fetal tissues of mice harbor bacterial communities using bacterial culture, qPCR, and 16S rRNA gene sequencing and by comparing the bacterial profiles of these tissues to those of maternal tissues and background technical controls.

## MATERIALS AND METHODS

### Study subjects and sample collection

C57BL/6 mice were purchased from The Jackson Laboratory in Bar Harbor, ME, USA, and bred in the animal care facility at C.S. Mott Center for Human Growth and Development at Wayne State University, Detroit, MI. Eleven pregnant mice were euthanized at 17.5 days of gestation. The dam’s chest and abdomen were shaved, 70% alcohol was applied, and the dam was placed on a surgical platform within a biological safety cabinet. Study personnel donned sterile surgical gowns, masks, full hoods, and powder-free exam gloves during sample collection. Sterile disposable scissors and forceps were used throughout, and new scissors and forceps were used for each organ that was sampled.

The oral cavity and vagina were swabbed with Dacron (Medical Packaging Corp., Camarillo, CA) and ESwabs (BD Diagnostics, Sparks, MD) for molecular microbiology and bacterial culture, respectively. For the abdomen, a Dacron swab was collected, iodine was applied and, after the iodine dried, an ESwab was collected. A midline incision was made along the full length of the abdomen. The peritoneum was sampled with a Dacron swab. The uterine horns were separated from the cervix and placed within a sterile Petri dish, wherein they were immediately processed by a different investigator within the biological safety cabinet. Uterine horns were dissected and fetuses (the fetus inside the amniotic sac attached to the placenta) were placed in individual Petri dishes. Uterine tissues were collected for both molecular microbiology and bacterial culture. Two fetuses from each dam were selected for analysis; tissues from one were used for molecular microbiology and tissues from the other for bacterial culture. From each fetus, the placenta, lung, liver, intestine, and brain were collected (molecular microbiology was performed on fetal brain samples from all 11 mice, i.e. mice A-K; bacterial culture was completed on fetal brain samples from mice E-K). The fetal spleen and tail were also collected for molecular microbiology.

Next, the maternal cervix, liver, and lung were sectioned and one sample of each was placed into a sterile 1.5 ml microcentrifuge tube and an anaerobic transport medium tube (Anaerobe Systems, Morgan Hill, CA) for molecular microbiology and bacterial culture, respectively. Lastly, after all placental and fetal tissues were sampled and stored, the maternal heart and the maternal proximal and distal large intestine were collected for molecular microbiology, and the maternal middle intestine was collected for bacterial culture. Procedures were approved by the Institutional Animal Care and Use Committee at Wayne State University (Protocol 18-03-0584).

### Bacterial culture

ESwabs and tissues collected for bacterial culture were placed within a COY Laboratory Products (Grass Lake, MI) hypoxic growth chamber (5% CO_2_, 5% O_2_) and processed in the following order: placenta, fetal liver, fetal lung, fetal brain, fetal intestine, maternal uterus, maternal liver, maternal lung, maternal cervix, maternal skin post-sterilization, maternal vagina, maternal oral cavity, and the maternal mid-intestine. While processing samples for bacterial culture within the chamber, study personnel wore sterile sleeve protectors (Kimtech pure A5; 36077; Kimberly Clark, Irving, TX), nitrile exam gloves, and sterile nitrile gloves (52102; Kimberly Clark) over the top of the nitrile exam gloves.

Tissues were removed from the anaerobic transport medium tubes using a sterile disposable inoculating loop (10 µl; Fisher Scientific, Hampton, NH), placed into a sterilized Wheaton dounce reservoir (2 ml or 5 ml; DWK Life Sciences, Millville, NJ) containing 1 ml of sterile phosphate-buffered saline (PBS; Gibco, Fisher Scientific), and homogenized for 1 minute. Tissue homogenates were transferred to a 5 ml centrifuge tube containing an additional 1.5 ml of sterile PBS. For mice E-K, maternal lung and maternal mid-intestine tissues were homogenized in sterile 5 ml centrifuge tubes using 0.5 ml sterile PBS and a sterile disposable scalpel (Surgical Design, Lorton, VA).

Tissue homogenates and ESwab buffer solutions were plated on blood agar (TSA with 5% sheep blood) and chocolate agar, and incubated at 37° C under oxic, hypoxic (5% CO_2_, 5% O_2_), and anoxic (5% CO_2_, 10% H, 85% N) atmospheres. All samples were additionally plated on MacConkey agar, and also added to SP4 broth with urea and SP4 broth with arginine, and incubated at 37° C under an oxic atmosphere. All samples were cultured in duplicate under all growth conditions (media type x atmosphere) and were incubated for seven days. There was ultimately no growth of *Ureaplasma* or *Mycoplasma* spp. from maternal, placental, or fetal samples in SP4 broth, or of bacteria in general from placental and fetal samples on MacConkey agar. Therefore, data from these growth media (i.e. SP4 urea, SP4 arginine, MacConkey) are not included in the Results section.

During the processing of each mouse’s samples for culture, three chocolate agar plates were left open in the hypoxic chamber to serve as negative controls – they were subsequently incubated for seven days under oxic, hypoxic, and anoxic conditions. Additionally, for each mouse, the PBS stock used for tissue homogenization was plated on blood agar, chocolate agar and MacConkey agar, and was further added to SP4 broths containing urea and arginine. The PBS-control blood agar and chocolate agar plates were incubated under oxic, hypoxic, and anoxic conditions, and the MacConkey agar plates and SP4 broths were incubated under oxic conditions. These negative controls were incubated for seven days.

Distinct bacterial isolates (i.e. colonies) recovered from negative controls and placental or fetal tissues were streaked for purity and taxonomically identified based upon their 16S rRNA gene sequence identity, as determined through Sanger sequencing (see details below). In one instance, there was contiguous growth of bacterial isolates on a plate for a single placental sample (that of Mouse J); the isolates all had the same morphotype, so a representative isolate was streaked for purity and taxonomically identified through Sanger sequencing.

The negative control plates yielded five total bacterial isolates over the course of the experiment. Four were successfully sequenced: two were identified as *Cutibacterium acnes* and two as *Staphylococcus hominis*. If a specific bacterium was cultured on a technical control plate on the day a mouse’s samples were processed as well as on a placental or fetal sample plate for that mouse (i.e. there was a 100% 16S rRNA gene sequence match between the bacterial isolates recovered on the two plates), that bacterium was not included in analyses. Overall, this included 11 bacterial isolates for three mice (D, J & K). Of these 11 isolates, four were *Cutibacterium acnes* and seven were *Staphylococcus hominis*. If a specific bacterium was cultured on a mouse’s placental or fetal sample plate as well as on a technical control plate from another sample processing day, but not on a control plate from that mouse’s sampling day, the bacterium was included in analyses.

For maternal cervix, uterus, and liver samples, the unique isolate morphotypes on each plate were streaked for purity and taxonomically identified through Sanger sequencing of the 16S rRNA gene. Samples of the maternal oral cavity, lung, vagina, and intestine often yielded bacterial isolates with contiguous growth. Therefore, the taxonomic identities of the bacteria cultured from these samples were determined through plate wash PCR (46) followed by 16S rRNA gene sequencing (see details below).

During each week of the experiment, blood, chocolate, and MacConkey agar plates were inoculated with *Eikenella corrodens, Enterococcus faecalis, Escherichia coli, Klebsiella pneumoniae, Staphylococcus aureus*, and *Streptococcus agalactiae*, and cultured under oxic, hypoxic, and anoxic conditions. SP4 broths with urea or arginine were inoculated with *Ureaplasma urealyticum* and *Mycoplasma hominis*, respectively. Each of these cultures was positive over the course of the experiment (MacConkey agar was positive for *Escherichia coli* throughout).

### Taxonomic identification of individual bacterial isolates

After being streaked for purity, bacterial isolates from placental, fetal, and maternal uterine, cervical, and liver samples were stored in nuclease-free water and frozen at −20° C until colony PCR targeting the 16S rRNA gene was performed. The 16S rRNA gene of each isolate was first amplified using the 27F/1492R primer set and then bi-directionally Sanger sequenced through GENEWIZ (South Plainfield, NJ) using the 515F/806R primer set, which targets the V4 hypervariable region of the 16S rRNA gene. Forward and reverse reads were trimmed using DNA Baser software (http://www.dnabaser.com/) with default settings, and assembled using the CAP (contig assembly program) of BioEdit software (v7.0.5.3), also with default settings. The taxonomic identities of individual bacterial isolates were determined using the Basic Local Alignment Search Tool (BLAST) (47). 16S rRNA gene sequence similarities between isolates and their top match on BLAST were ≥ 99.5%, unless otherwise noted **(Table 1; Table 2)**.

**Table 1.** Bacterial cultivation results for placental and fetal brain, lung, liver, and intestinal samples.

| Mouse | Placental or fetal body site | Bacterial culture | | 16S rRNA gene qPCR | 16S rRNA gene sequence match between the isolate and $\geq 1$ sequence within a 16S rRNA gene library | |
| --- | --- | --- | --- | --- | --- | --- |
| | | Total # of isolates recovered | Top NCBI BLAST taxonomic designation ($\geq 99.5\%$ 16S rRNA gene sequence identity unless otherwise indicated) | Was sample bacterial load > that of blank kit controls? | Library for that specific tissue type in that Mouse | Library for any maternal body site in that Mouse |
| A | Placenta | 0 |  | No |  |  |
|  | Lung | 0 |  | No |  |  |
|  | Liver | 0 |  | No |  |  |
|  | Intestine | 0 |  | Yes |  |  |
| B | Placenta | 1 | *Cutibacterium acnes* | No | No | No |
|  | Lung | 0 |  | No |  |  |
|  | Liver | 1 | *Cutibacterium acnes* | No | No | No |
|  | Intestine | 0 |  | No |  |  |
| C | Placenta | 0 |  | No |  |  |
|  | Lung | 1 | *Bacillus simplex* / *frigoritolerans* | No | No | No |
|  | Liver | 0 |  | No |  |  |
|  | Intestine | 0 |  | No |  |  |
| D | Placenta | 0 |  | No |  |  |
|  | Lung | 0 |  | No |  |  |
|  | Liver | 0 |  | No |  |  |
|  | Intestine | 0 |  | No |  |  |
| E | Placenta | 1 | *Corynebacterium tuberculostearicum* (98.5%) | Yes | No | No |
|  | Brain | 1 | *Bacillus halosaccharovorans* | Yes | No | No |
|  | Lung | 0 |  | No |  |  |
|  | Liver | 0 |  | No |  |  |
|  | Intestine | 0 |  | No |  |  |
| F | Placenta | 1 | *Staphylococcus hominis* | No | No | Yes (lung, skin) |
|  | Brain | 7 | *Bacillus circulans*; *Bacillus megaterium* / *flexus*; *Bacillus* spp.; *Ornithinibacillus* sp. Marseille-P3601; *Paenibacillus* spp. | Yes | No | Yes, for 1/7 isolates (skin) |
|  | Lung | 0 |  | No |  |  |
|  | Liver | 1 | *Bacillus sonorensis* | No | No | No |
|  | Intestine | 1 | *Staphylococcus hominis* (99.4%) | No | No | Yes (lung, skin) |
| G | Placenta | 0 |  | No |  |  |
|  | Brain | 1 | *Staphylococcus hominis* | No | No | Yes (peritoneum, skin) |
|  | Lung | 0 |  | No |  |  |
|  | Liver | 1 | *Staphylococcus hominis* | No | No | Yes (peritoneum, skin) |
|  | Intestine | 0 |  | No |  |  |
| H | Placenta | 16 | *Staphylococcus hominis*; *Staphylococcus epidermidis* / *caprae* / *capitis* | No | No | No |
|  | Brain | 3 | *Staphylococcus hominis*; *Staphylococcus warneri*; *Staphylococcus epidermidis* / *caprae* / *capitis* | No | No | No |
|  | Lung | 0 |  | No |  |  |
|  | Liver | 1 | *Paenibacillus timonensis* (98.0%) | No | No | No |
|  | Intestine | 0 |  | Yes |  |  |
| I | Placenta | 0 |  | No |  |  |
|  | Brain | 0 |  | No |  |  |
|  | Lung | 1 | *Cutibacterium acnes* (99.0%) | No | No | No |
|  | Liver | 0 |  | No |  |  |
|  | Intestine | 0 |  | No |  |  |
| J | Placenta | TMTC* | *Staphylococcus caprae* | No | Yes | Yes (heart, mouth, intestine) |
|  | Brain | 0 |  | No |  |  |
|  | Lung | 0 |  | No |  |  |
|  | Liver | 0 |  | No |  |  |
|  | Intestine | 0 |  | No |  |  |
| K | Placenta | 0 |  | No |  |  |
|  | Brain | 0 |  | No |  |  |
|  | Lung | 0 |  | No |  |  |
|  | Liver | 0 |  | No |  |  |
|  | Intestine | 0 |  | No |  |  |

**Table 2.** Bacterial cultivation results for maternal cervical, uterine, and liver samples.

| | | Bacterial culture | | 16S rRNA gene qPCR | 16S rRNA gene sequence match between the isolate and $\geq 1$ sequence within a 16S rRNA gene library |
| --- | --- | --- | --- | --- | --- |
| Mouse | Low microbial biomass maternal body site | # of unique colony morphotypes recovered | Top NCBI BLAST taxonomic designation ($\geq 99.5\%$ 16S rRNA gene sequence identity unless otherwise indicated) | Was sample bacterial load > that of blank kit controls? | Library for that specific tissue type in that Mouse |
| A | Cervix | 1 | *Rodentibacter pneumotropicus* (98.0%) | No | Yes |
|  | Uterus | 1 | *Bacillus niabensis* | No | No |
|  | Liver | 0 |  | No |  |
| B | Cervix | 0 |  | No |  |
|  | Uterus | 0 |  | No |  |
|  | Liver | 2 | *Lactobacillus gasseri*; *L. murinus* | No | Yes, for 1/2 morphotypes |
| C | Cervix | 0 |  | No |  |
|  | Uterus | 0 |  | No |  |
|  | Liver | 5 | *Bacteroides sartorii* (98.0%); *Klebsiella variicola*; *L. gasseri* / *L. johnsonii*; *L. murinus*; *Staphylococcus hominis* | Yes | Yes, for 2/5 morphotypes |
| D | Cervix | 0 |  | No |  |
|  | Uterus | 0 |  | No |  |
|  | Liver | 0 |  | No |  |
| E | Cervix | 1 | *Pasteurella caecimuris* | Yes | Yes |
|  | Uterus | 0 |  | No |  |
|  | Liver | 0 |  | No |  |
| F | Cervix | 0 |  | Yes |  |
|  | Uterus | 0 |  | No |  |
|  | Liver | 1 | *Staphylococcus epidermidis* / *S. caprae* / *S. capitis* | No | No |
| G | Cervix | 5 | *Bacteroides sartorii*; *Faecalibaculum rodentium* (97.9%); *Lactobacillus murinus*; *L. reuteri* (99.2%); *Pasteurella caecimuris* | Yes | Yes, for 5/5 morphotypes |
|  | Uterus | 0 |  | No |  |
|  | Liver | 0 |  | No |  |
| H | Cervix | 0 |  | Yes |  |
|  | Uterus | 0 |  | No |  |
|  | Liver | 0 |  | No |  |
| I | Cervix | 6 | *Bacillus circulans*; *Pasteurella caecimuris*; *Rodentibacter pneumotropicus* (98.1%); *Staphylococcus hominis*; *S. xylosus*; *Streptococcus thoraltensis* (99.3%) | No | Yes, for 4/6 morphotypes |
|  | Uterus | 0 |  | No |  |
|  | Liver | 0 |  | No |  |
| J | Cervix | 1 | *Pasteurella caecimuris* | No | Yes |
|  | Uterus | 1 | *Staphylococcus aureus* | No | No |
|  | Liver | 1 | *Staphylococcus epidermidis* / *S. caprae* / *S. capitis* | No | No |
| K | Cervix | 1 | *Pasteurella caecimuris* | Yes | Yes |
|  | Uterus | 0 |  | Yes |  |
|  | Liver | 0 |  | No |  |

### DNA extraction from plate washes of cultured bacteria

Plate wash was performed by pipetting 1-2 ml of PBS onto the agar plate and dislodging bacterial colonies with either sterile L-shaped spreaders or inoculating loops. The PBS wash was then transferred into cryovials and stored at −80°C until DNA was extracted. DNA was extracted from plate wash samples using Qiagen DNeasy PowerSoil (Germantown, MD) extraction kits. Washes from maternal samples that yielded growth under multiple atmospheres for the same media type were pooled prior to the extraction process. Purified DNA was stored at −20° C.

### 16S rRNA gene sequencing of plate wash extracts

The 16S rRNA genes in plate wash extracts were sequenced at Wayne State University on an Illumina MiSeq system using a 2 × 250 cycle V2 kit, and following Illumina sequencing protocols (48). The 515F/806R primer set was used to target the V4 region of the 16S rRNA gene. The 16S rRNA gene sequences from the paired fastq files for these samples were processed as previously described (21).

### DNA extraction from swab and tissue samples

All Dacron swab and tissue samples collected for molecular microbiology were stored at - 80° C until DNA extractions were performed. DNA extractions were performed in a biological safety cabinet by study personnel donning sterile surgical gowns, masks, full hoods, and powder-free exam gloves. Extractions of tissues generally included 0.015 – 0.100 grams of tissue, except for the fetal tail and spleen, whose masses were very low.

DNA was extracted from swabs, tissues, and background technical controls (i.e. sterile Dacron swabs (N = 11) and blank DNA extraction kits (N = 23)) using the DNeasy PowerLyzer PowerSoil Kit (Qiagen, Germantown, MD) with minor modifications to the manufacturer’s protocol. Specifically, 400 μl of bead solution, 200 μl of phenol:chloroform:isoamyl alcohol (pH 7–8), and 60 μl of Solution C1 were added to the supplied bead tube. Cells within samples were lysed by mechanical disruption for 30 seconds using a bead beater. After centrifugation, the supernatants were transferred to new tubes, and 100 μl of solution C2, 100 μl of solution C3, and one μl of RNase A enzyme were added, and tubes were incubated at 4° C for five minutes. After centrifugation, the supernatants were transferred to new tubes that contained 650 μl of solution C4 and 650 μl of 100% ethanol. The lysates were loaded onto filter columns, centrifuged for one minute, and the flow-through was discarded. This step was repeated until all sample lysates were spun through the filter columns. Five hundred μl of solution C5 were added to the filter columns, centrifuged for one minute, the flow-through was discarded, and the tube was centrifuged for an additional three minutes as a dry-spin. Finally, 60 μl of solution C6 were placed on the filter column and incubated for five minutes before centrifuging for 30 seconds to elute the extracted DNA. Purified DNA was stored at −20° C.

Purified DNA was quantified using a Qubit 3.0 fluorometer with a Qubit dsDNA BR Assay kit (Life Technologies, Carlsbad, CA), according to the manufacturer’s protocol. All purified DNA samples were then normalized to 80 ng/µl (when possible) by diluting each sample with the Qiagen elution buffer (Solution C6).

### 16S rRNA gene quantitative real-time PCR (qPCR)

A preliminary test was performed to investigate whether DNA amplification inhibition existed among the different sample types. For this test, 4.7 μl of purified *Escherichia coli* ATCC 25922 (GenBank accession: CP009072) genomic DNA (0.005 ng/µl) containing seven 16S rDNA copies per genome was spiked into 7.0 μl of purified DNA from mouse samples that were serially diluted with Solution C6 by a factor of 1:3 (i.e. 1:0, 1:3, 1:9). For tissue sample types with a mean DNA concentration above 250 ng/µl, DNA concentrations were normalized to 80 ng/μl by dilution with Solution C6 before being serially diluted and spiked with *E. coli* genomic DNA. Genomic DNA was quantified using a Qubit 3.0 fluorometer with a Qubit dsDNA HS Assay kit (Life Technologies, Carlsbad, CA) according to the manufacturer’s protocol. Three μl of each spiked sample were then used as a template for qPCR. For all samples, spiked reactions contained approximately 1.0 × 10^3^ *E. coli* 16S rDNA copies. There was no evidence of DNA amplification inhibition **(Supplemental Figure 1A, B)**.

Total bacterial DNA abundance within samples was measured via amplification of the V1 - V2 region of the 16S rRNA gene according to the protocol of Dickson et al (49) with minor modifications. These modifications included the use of a degenerative forward primer (27f-CM: 5’-AGA GTT TGA TCM TGG CTC AG-3’) (50) and a degenerate probe containing locked nucleic acids (+) (BSR65/17: 5’-56FAM-TAA +YA+C ATG +CA+A GT+C GA-BHQ1-3’). Each 20 μl reaction contained 0.6 μM of 27f-CM primer, 0.6 μM of 357R primer (5’-CTG CTG CCT YCC GTA G-3’), 0.25 μM of BSR65/17 probe, 10.0 μl of 2X TaqMan Environmental Master Mix 2.0 (Life Technologies, Carlsbad, CA), and 3.0 μl of either purified DNA (diluted to 80 ng/µl when possible), elution buffer, or nuclease-free water. The total bacterial DNA qPCR was performed using the following conditions: 95° C for 10 min, followed by 45 cycles of 94° C for 30 sec, 50° C for 30 sec, and 72° C for 30 sec. Duplicate reactions were run for all samples. All samples were run across a total of five runs.

Raw amplification data were normalized to the ROX passive reference dye and analyzed using the on-line platform Thermo Fisher Cloud: Standard Curve (SR) 3.3.0-SR2-build15 with automatic threshold and baseline settings. Cycle of quantification (Cq) values were calculated for samples based on the mean number of cycles required for normalized fluorescence to exponentially increase.

After plotting a regression of log(*E. coli* 16S rRNA gene copy number) and Cq value for standard curves included in each qPCR run, 16S rRNA gene copy number in mouse samples was calculated according to Gallup (51) using the equation *X*_*o*_ = *E*_*AMP*_^(*b*-*Cq*)^, where *E*_*AMP*_ is the exponential amplification value for the qPCR assay, calculated as *E*_*AMP*_ = 10^(−1/*m*)^ and *b* and *m* are the intercept and slope of the regression.

### 16S rRNA gene sequencing of swab and tissue sample extracts

Amplification and sequencing of the V4 region of the 16S rRNA gene was performed at the University of Michigan’s Center for Microbial Systems as previously described (52), except that library builds were performed in triplicate and pooled for each individual sample prior to the equimolar pooling of all sample libraries for multiplex sequencing. Sample-specific MiSeq run files have been deposited on the NCBI Sequence Read Archive (BioProject ID SUB6641162).

Raw sequence reads were processed using mothur software (v1.39.5) (53) following the Standard Operating Procedure provided by Schloss et al. (www.mothur.org/wiki/MiSeq_SOP). Paired-end reads were assembled into contiguous sequences, quality checked (maximum length = 275, maximum ambiguous base pairs = 0, and maximum number of homopolymers = 8), and aligned against the SILVA 16S rDNA reference database (release 102) (54, 55); sequences falling outside the target alignment space were removed. Quality sequences were pre-clustered (diffs = 2) and chimeric sequences were identified with VSEARCH (56) and removed. The remaining sequences were taxonomically classified using the SILVA reference database with a k-nearest neighbor approach and a confidence threshold of 80%. Sequences derived from an unknown domain, Eukaryota, Archaea, chloroplasts, or mitochondria were removed. Operational taxonomic units (OTUs) were defined by clustering sequences at a 97% sequence similarity cutoff using the average neighbor method.

### Statistical analysis

The bacterial loads, as assessed through qPCR, of maternal, placental and fetal samples were compared to those of background technical controls (i.e. sterile Dacron swabs and blank DNA extraction kits) using t-tests or Mann-Whitney U tests with sequential Bonferroni corrections applied. The bacterial loads of placental and fetal tissues were compared to one another using Wilcoxon matched pairs tests, again corrected for multiple comparisons.

The beta diversity of 16S rRNA gene profiles among maternal, placental, fetal and technical control samples were characterized using the Bray-Curtis similarity index. Bray-Curtis similarities in sample profiles were visualized using Principal Coordinates Analysis (PCoA) plots and statistically evaluated using non-parametric multivariate ANOVA (NPMANOVA). These analyses were limited to samples that yielded a 16S rRNA gene library with ≥ 250 quality-filtered sequences and a Good’s coverage ≥ 95%. All data analysis was completed in PAST software (v 3.25) (57). Heat maps of sample bacterial profiles were generated using the open-source software program Morpheus (https://software.broadinstitute.org/morpheus).

## RESULTS

### Bacterial culture from placental and fetal tissues

Growth of bacterial isolates from placental and fetal tissues was rare **(Figure 1A; Figure 2).** Only 3/11 mice (F, H & J) yielded more than two total bacterial isolates across all their cultured placental and fetal samples under all growth conditions **(Table 1)**. Most of the bacterial isolates from placental and fetal samples were *Staphylococcus* spp. (mostly *S. hominis*) **(Figure 1A)**. *Staphylococcus* spp. were cultured from the mouth, intestine, and vagina of dams **(Figure 1B)**; however, two of the five bacterial isolates recovered from the 114 negative control plates included in this study were also *Staphylococcus*, specifically *S. hominis*. The non-staphylococci bacteria cultured from placental or fetal samples were *Bacillus, Corynebacterium, Paenibacillus, Propionibacterium*, and unclassified bacilli **(Table 1)**. These bacteria were rarely, if ever, cultured from maternal samples **(Figure 1A, B)**.

**Figure 1.**
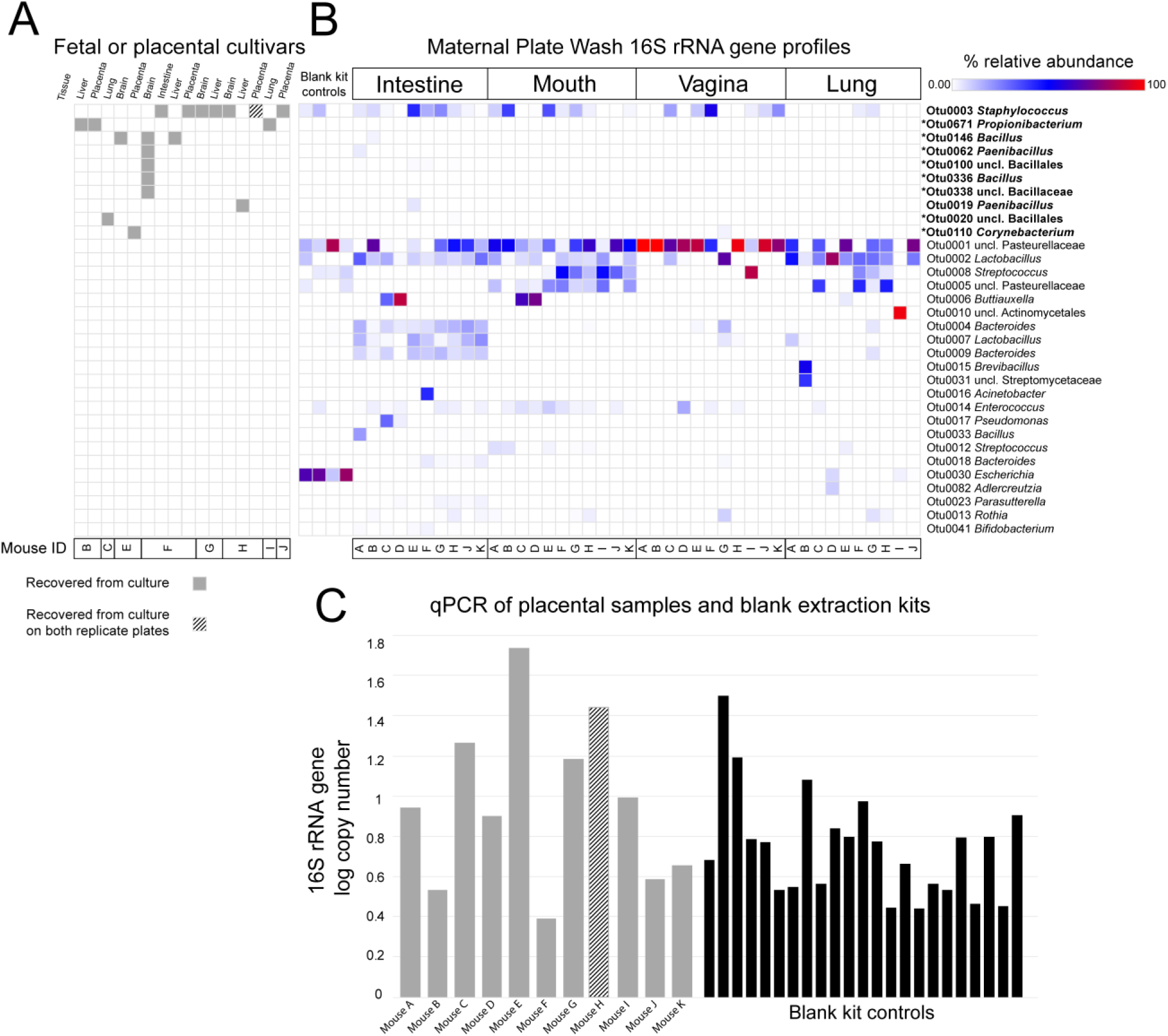
Bacterial cultivation results for A) fetal and placental tissues in relation to those for B) maternal intestinal, mouth, vaginal, and lung samples, and C) a comparison of the bacterial loads of individual placental samples and blank extraction kit controls in light of the cultivation results. Panel A indicates the recovery of bacterial isolates from placenta and/or fetal tissues, by mouse and across different growth media and atmosphere conditions. The taxonomic assignments of these isolates were determined by comparing their 16S rRNA gene sequences to those of the operational taxonomic units (OTUs) of molecular surveys of the mixed bacterial communities cultured from maternal intestinal, oral, vaginal, and lung samples (sequence identity was ≥ 97.2%). Panel B provides the results of 16S rRNA gene molecular surveys of the plate washes of bacterial growth from maternal intestinal, oral, vaginal, and lung samples, as well as of blank extraction kit controls processed alongside the plate washes. OTUs were included in the heat map in Panel B if they had an average percent relative abundance ≥ 0.5% across all plate washes or if they were the best 16S rRNA gene sequence match to bacterial isolates in Panel A (indicated by an asterisk). The bolded OTUs represent the best 16S rRNA gene sequence matches to placental and fetal isolates in Panel A. Panel C illustrates similarities in bacterial load, as assessed by 16S rRNA gene quantitative real-time PCR (qPCR), between placental samples yielding at least one bacterial isolate and blank DNA extraction kit controls.

**Figure 2.**
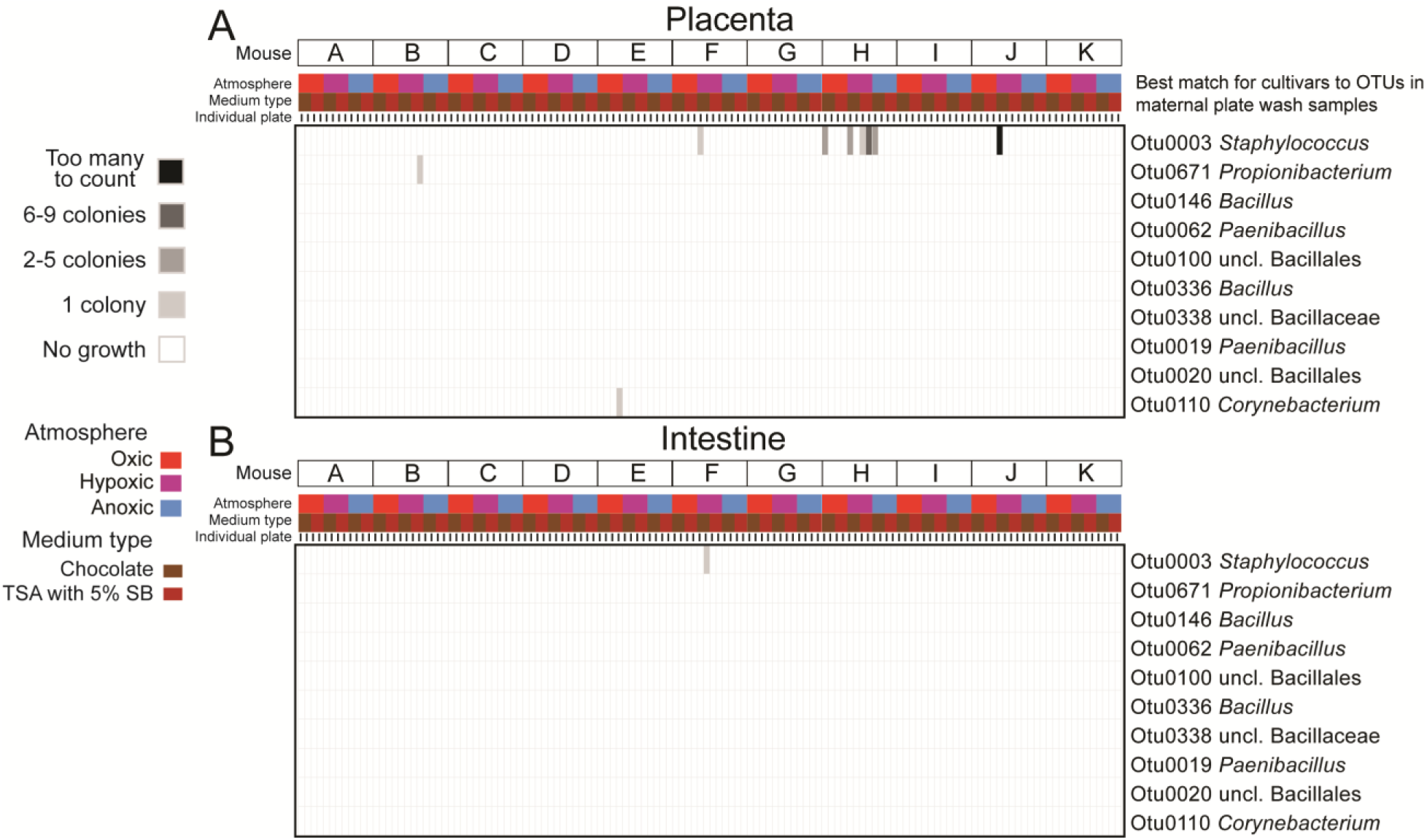
Heat maps illustrating bacterial cultivation results for A) placenta and B) fetal intestinal tissues. Each column of the heat map represents a single agar plate. The x-axis indicates the mouse identity, atmospheric condition, growth medium, and paired replicate for each agar plate. The vast majority of blood and chocolate agar plates did not yield any bacterial growth over seven days for placental (93.2%) and fetal intestinal (99.2%) samples. The operational taxonomic units (OTUs) on the y-axis are those that represent the best 16S rRNA gene sequence matches to bacterial isolates recovered from any placental or fetal sample in this study overall (i.e. the OTUs in bold font in Figure 1B).

In general, only one or two placental or fetal sites within a given fetus yielded a bacterial isolate, and there was little consistency among the fetuses in terms of which site yielded an isolate **(Figure 1A; Table 1)**. For example, of the 132 blood and chocolate agar plates on which placental tissue homogenates were spread, only nine (6.8%) yielded even a single bacterial isolate, and five of these plates came from a single placental sample (Mouse H) **(Figure 2)**. All of the bacterial isolates from Mouse H’s placental sample were *Staphylococcus* (either *S. hominis* or *S. epidermidis* / *caprae* / *capitis*). There were no exact matches of the 16S rRNA genes of these isolates within the 16S rRNA gene surveys of placental tissues from Mouse H, nor were there any matches within the 16S rRNA gene surveys of any of the sampled maternal body sites for Mouse H, which included the maternal skin, heart, mouth, lung, liver, proximal intestine, distal intestine, peritoneum, cervix, and vagina **(Table 1)**. The placental sample from Mouse J yielded many colonies of *Staphylococcus caprae* on one chocolate agar plate under hypoxic conditions; yet there were no bacterial colonies on the replicate chocolate agar plate incubated under hypoxic conditions, nor on any other plate for this sample **(Figure 2; Table 1)**. An exact match of the 16S rRNA gene of this *Staphylococcus caprae* isolate was identified in the 16S rRNA gene survey of placental tissues from Mouse J, as well as in the 16S rRNA gene surveys of the maternal heart, mouth, and proximal intestine samples for Mouse J. However, the bacterial load of the placental sample from Mouse J, as assessed by 16S rRNA gene qPCR, was not high – it was less than the bacterial load of 14/23 (60.9%) DNA extraction kit controls **(Figure 1C)**.

Of the 132 blood and chocolate agar plates on which fetal intestinal tissue homogenates were spread, only one yielded growth – a single bacterial colony of *Staphylococcus hominis* **(Figure 2)**. The 16S rRNA gene of this bacterial isolate was not detected in the molecular survey of fetal intestines from this mouse (Mouse F), but it was identified in the 16S rRNA gene surveys of maternal lung and skin from Mouse F **(Table 1)**. This sample had the lowest bacterial load of any fetal intestinal sample in the study, and had a bacterial load less than that of 14/23 (60.9%) DNA extraction kit controls **(Figure 3)**.

**Figure 3.**
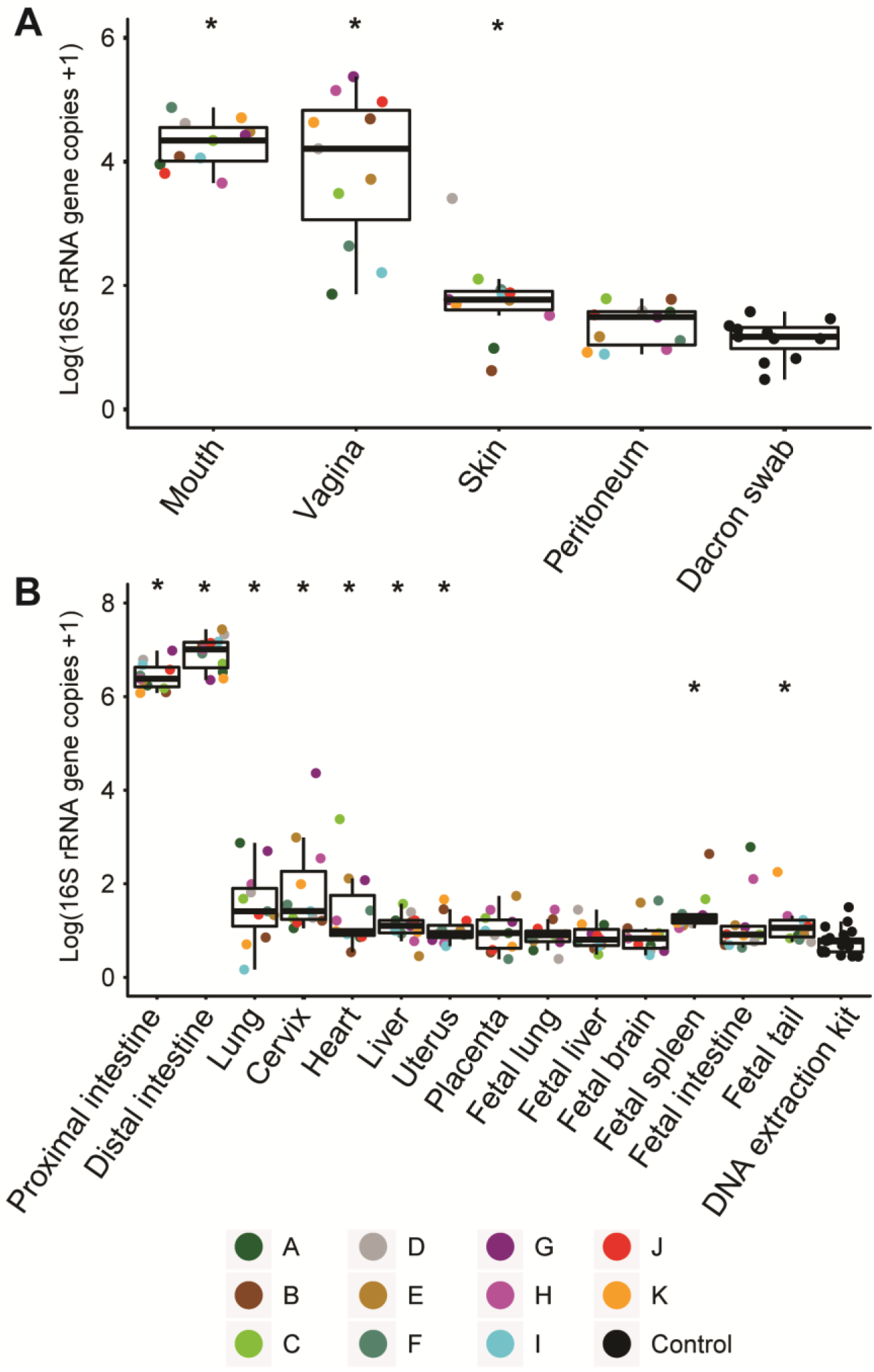
Quantitative real-time PCR (qPCR) analyses illustrating variation in bacterial load among A) maternal swab samples and Dacron swab controls, and B) maternal, placental, and fetal tissue samples and blank DNA extraction kit controls. Bars indicate the median and quartile log-16S rRNA gene copy values for each sample and control type. Points, color-coded by mouse identity, indicate the mean values of two replicate qPCR reactions. An asterisk indicates that bacterial loads of that sample type were greater than those of corresponding technical controls.

### Bacterial culture from maternal compartments

Bacterial cultures of the maternal intestine, mouth, vagina, and lung often yielded lawns of bacterial growth dominated by unclassified Pasteurellaceae, *Lactobacillus*, and *Staphylococcus* **(Figure 1B)**. Body site-specific variation in the structure of cultured bacterial communities from maternal samples was evident **(Figure 1B)**. For instance, the vast majority of bacteria cultured from the vagina were unclassified Pasteurellaceae, while *Bacteroides* and a distinct strain of *Lactobacillus* were consistently cultured from the maternal intestine in addition to the unclassified Pasteurellaceae, *Lactobacillus*, and *Staphylococcus* isolated from other body sites **(Figure 1B)**.

Bacterial cultures of the maternal cervix yielded isolates in 6/11 (54.5%) mice **(Table 2)**. The most common bacterium cultured from the murine cervix was *Pasteurella caecimuris*; it was recovered in culture from 5/11 cervical samples. In each case, an exact match for the 16S rRNA gene of the *Pasteurella caecimuris* isolate was identified in the 16S rRNA gene survey of the corresponding cervical sample **(Table 2)**.

Bacteria were rarely cultured from the uterus (2/11 mice) and maternal liver (4/11 mice) **(Table 2)**. The two bacteria cultured from the uterus were *Bacillus niabensis* and *Staphylococcus aureus*. An exact match of the 16S rRNA gene of these isolates was not identified in the 16S rRNA gene surveys of the respective uterine samples. The bacteria cultured from maternal liver samples were primarily *Lactobacillus* and *Staphylococcus* species. Of the nine distinct bacterial morphotypes cultured from maternal liver tissues, only 3 (33%) had an exact match of their 16S rRNA gene identified in the 16S rRNA gene surveys of their respective samples **(Table 2)**.

### Quantitative real-time PCR (qPCR) of murine and control samples

Bacterial load, as characterized by 16S rRNA gene copy abundance, varied greatly across maternal, placental, and fetal body sites **(Figure 3)**. The bacterial loads of swabs of the maternal mouth, vagina, and skin exceeded those of sterile Dacron swabs **(Figure 3A)**. Similarly, the bacterial loads of tissues of the maternal proximal and distal intestine, lung, cervix, heart, liver, and uterus exceeded those of blank DNA extraction kits **(Figure 3B)**. In contrast, bacterial loads of the maternal peritoneum, the placenta, and the fetal lung, liver, brain, and intestine did not exceed those of their respective background technical controls **(Figure 3A, B)**. The spleen and tail were the only fetal tissues with bacterial loads exceeding those of blank DNA extraction kits **(Figure 3B)**. However, only 1/11 (9.1%) fetal tail and 2/11 (18.2%) fetal spleen samples had bacterial loads exceeding those of each of the blank DNA extraction kits. Corrected for multiple comparisons, no placental or fetal tissue, including the tail and spleen, had a bacterial load exceeding that of any other placental or fetal tissue (Wilcoxon matched pairs, p ≥ 0.68).

### 16S rRNA gene sequencing of murine and control samples

Six of the 23 (26.1%) blank DNA extraction kits, and 8/11 (72.7%) sterile swab controls, yielded a 16S rRNA gene library with ≥ 250 quality-filtered sequences and a Good’s coverage ≥ 95%. The prominent (i.e. ≥ 2.25% relative abundance) operational taxonomic units (OTUs) in the bacterial profiles of the DNA extraction kit controls were identified as *Ralstonia*, unclassified Bacillales, *Flavobacterium*, S24-7, *Brevibacterium, Pelomonas*, unclassified Bacteroidetes, and *Acinetobacter* **(Figure 4)**. However, only two of these prominent OTUs, identified as *Ralstonia* and *Pelomonas*, were present in the bacterial profiles of more than half of the DNA extraction kit controls. A decontam analysis indicated that the OTUs identified as *Ralstonia, Pelomonas, Pseudomonas*, and *Acinetobacter* were likely background DNA contaminants **(Figure 4)**.

**Figure 4.**
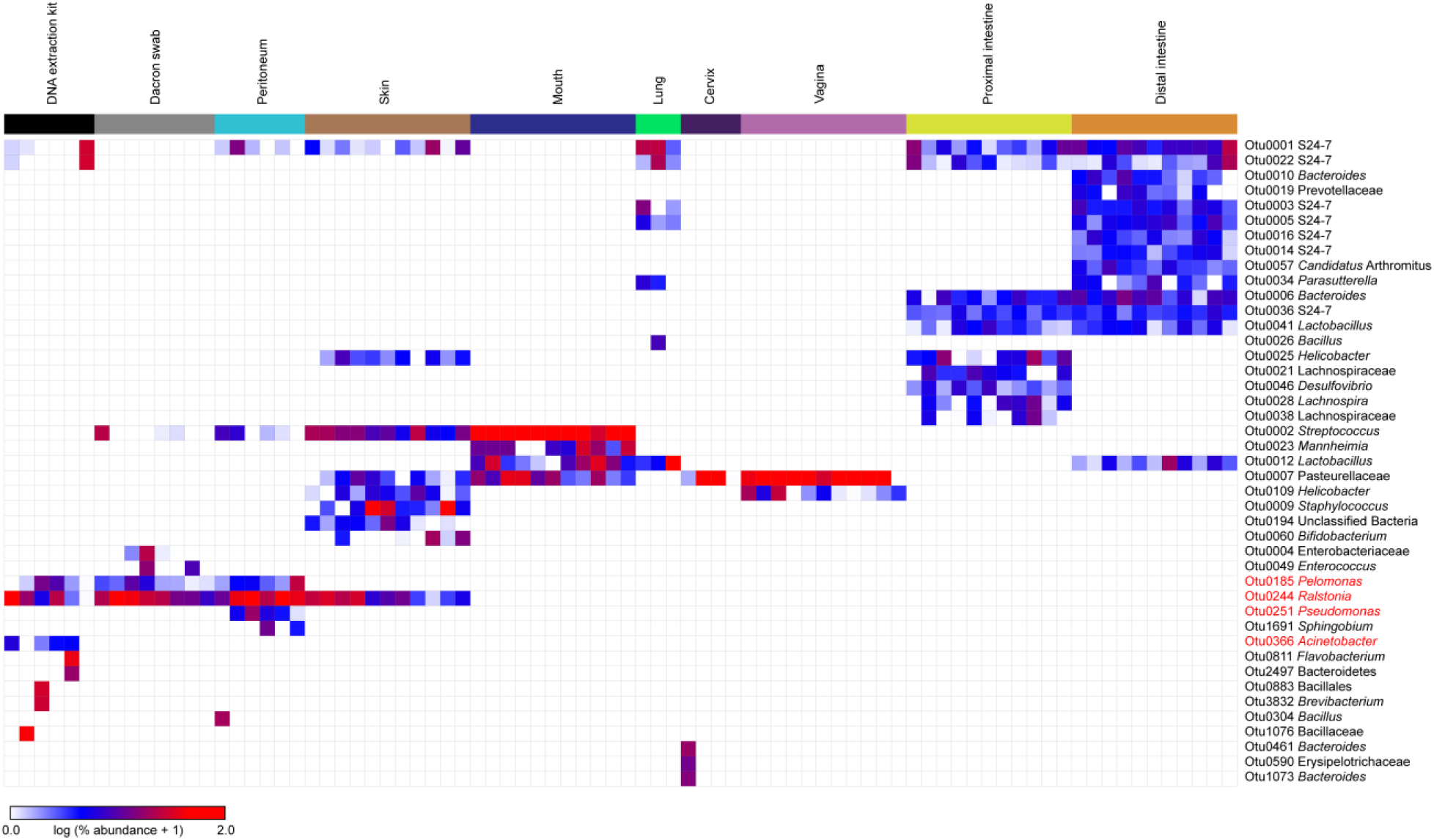
Heat map illustrating the relative abundances of prominent (≥ 2.25% average relative abundance) operational taxonomic units (OTUs) among the 16S rRNA gene profiles of maternal swab and tissue samples and background technical controls. The four OTUs in red font were identified as background DNA contaminants by the R package decontam.

The bacterial profiles of placental and fetal samples could not be compared to those of background technical controls because only two of the 77 (2.6%) placental and fetal brain, lung, liver, intestine, spleen, and tail samples included in this study, yielded a 16S rRNA gene library with ≥ 250 sequences and a Good’s coverage ≥ 95%. These two samples were the placenta from Mouse I and the fetal spleen from Mouse B. The placenta from Mouse I had an average bacterial load in comparison to that of other placentas **(Figure 3)**, and no bacteria were cultured from the placental tissues of this mouse **(Figure 1, Figure 2, Table 1)**. The prominent OTUs in the bacterial profile of the placental sample from Mouse I were identified as *Bacteroides, Akkermansia*, S24-7, *Lactobacillus*, and *Escherichia*. The fetal spleen from Mouse B had the highest bacterial load of any fetal spleen sample; its bacterial load was 58% higher than any other spleen sample **(Figure 3)**. The prominent OTUs in the bacterial profile of the fetal spleen from Mouse B were *Lactobacillus*, S24-7, and unclassified Lachnospiraceae.

All maternal skin, mouth, proximal and distal intestinal samples yielded a 16S rRNA gene library with ≥ 250 sequences and a Good’s coverage ≥ 95%. Six (54.5%), four (36.4%), and three (27.3%) maternal peritoneal, cervical, and lung samples, respectively, yielded a 16S rRNA gene library with ≥ 250 sequences and a Good’s coverage ≥ 95%. However, no maternal liver or uterine samples, and only one (9.1%) maternal heart sample, yielded a 16S rRNA gene library. The structure of the bacterial profiles of the maternal body sites with at least three 16S rRNA gene libraries meeting the above criteria were compared with those of background technical controls **(Figure 4, Figure 5)**.

**Figure 5.**
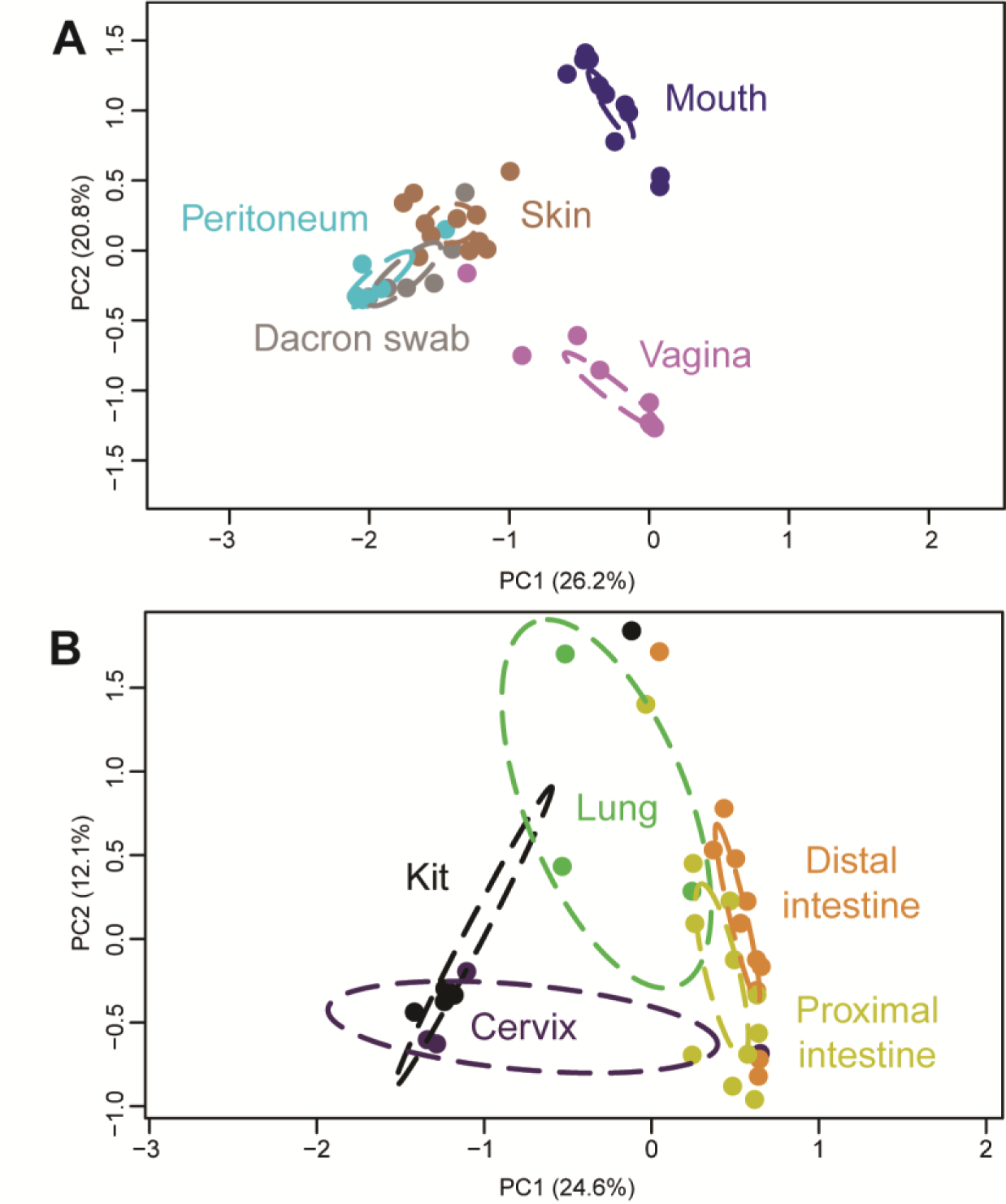
Principal Coordinates Analysis (PCoA) illustrating variation in 16S rRNA gene profiles among A) maternal swab samples and Dacron swab controls, and B) maternal tissue samples and blank DNA extraction kit controls. 16S rRNA gene profiles were characterized using the Bray-Curtis similarity index.

The taxonomic identities of prominent OTUs varied among maternal body sites **(Figure 4)**. Maternal proximal and distal intestinal samples had the most OTU-rich bacterial profiles. The maternal proximal intestine was characterized by *Bacteroides, Desulfovibrio, Helicobacter, Lachnospira*, unclassified Lachnospiraceae, *Lactobacillus*, and S24-7, while the maternal distal intestine had bacterial profiles consistently comprised of “*Candidatus* Arthromitus,” *Bacteroides, Lactobacillus, Parasutterella*, unclassified Prevotellaceae, and S24-7. Maternal vaginal and cervical bacterial profiles were dominated by unclassified Pasteurellaceae; the vagina also consistently contained *Helicobacter*. Maternal lung bacterial profiles were typified by *Lactobacillus* and S24-7, while those of the maternal mouth were dominated by *Streptococcus, Mannheimia, Lactobacillus*, and unclassified Pasteurellaceae. Maternal skin, a low microbial biomass site **(Figure 3A)**, and the peritoneum, a very low to nonexistent microbial biomass site **(Figure 3A)**, had bacterial profiles that overlapped with those of background technical controls more so than did the profiles of higher microbial biomass sites **(Figure 4)**. Specifically, skin bacterial profiles consistently contained *Bifidobacterium, Helicobacter*, unclassified Pasteurellaceae, *Ralstonia*, S24-7, *Staphylococcus*, and *Streptococcus. Ralstonia* was the dominant OTU in the bacterial profiles of the maternal peritoneum, as well as in the profiles of the background technical controls **(Figure 4)**. Indeed, the bacterial profiles of the maternal peritoneum were not distinguishable from those of background technical controls (Bray-Curtis similarity index; NPMANOVA, F = 0.974, p = 0.467) **(Figure 5)**.

### Comprehensive consideration of individual placental and fetal tissues across microbiological inquiries

Overall, there was only a single bacterial isolate (*Bacillus circulans*, cultured from the fetal brain tissue of Mouse F) that was cultured from a placental or fetal tissue that had a bacterial load higher than that of background technical controls, and that was identified in the 16S rRNA gene surveys of at least one of that fetus’ maternal samples **(Table 1)**.

## DISCUSSION

### Principal findings of the study

1) Of the 165 total bacterial cultures of placentas from the 11 mice, only nine (5.5%) yielded even a single colony, and five of those nine positive cultures came from a single mouse; 2) of the 165 total bacterial cultures of fetal intestinal tissues, only one (0.6%) was positive, yielding a single isolate of *Staphylococcus hominis*; 3) the bacterial loads of placental and fetal brain, lung, liver, and intestinal samples were not higher than those of DNA extraction kit controls; 4) only two (2.6%) placental or fetal tissue samples yielded a 16S rRNA gene library with at least 250 sequences and a Good’s coverage value of 95%; 5) the 16S rRNA gene libraries of each maternal skin, mouth, vaginal, and proximal and distal intestinal sample met these criteria, as did at least 25% of maternal lung, cervical, and peritoneum samples; 6) similar to the placental and fetal tissues samples, maternal heart, liver, and uterine samples did not yield 16S rRNA gene libraries with at least 250 sequences and a Good’s coverage value of 95%; and 7) overall, from all placental or fetal tissues for which there were culture, qPCR, and corresponding maternal sample sequence data (N = 49), there was only a single bacterial isolate that came from a fetal brain sample having a bacterial load higher than that of contamination controls and that was identified in sequence-based surveys of at least one of its corresponding maternal samples.

### Prior reports of placental and fetal microbiota in mice

An initial investigation of the existence of microbiota in the murine placenta and fetal intestine was carried out by Martinez et al. (40). Specifically, bacterial culture, 16S rRNA gene qPCR, and 16S rRNA gene sequencing were performed on the placenta and fetal intestines of 13 mice at day 17 of gestation (40). All bacterial cultures of the placenta and fetal intestine were negative. Yet, the bacterial loads of the fetal intestine were higher than those of placentas. After removing the OTUs detected in 16S rRNA gene sequencing surveys of background control samples from the overall dataset, the bacterial profiles of murine fetal intestine were dominated by *Enterococcus, Stramenopiles, Rhodoplanes*, and *Novosphingobium*. In contrast, the bacterial profiles of murine placentas were more diverse, with Pirellulaceae, Aeromonadaceae, MIZ46, ZB2, Veillonellaceae, Weeksellaceae, *Fluviicola, Bdellovibrio*, and Comamonadaceae being most common. The conclusion of the study was that, although murine fetuses do not appear to be populated by microbial communities, they are exposed to bacterial DNA *in utero*. Conversely, in a subsequent molecular study by Kuperman et al. (18), 24 murine placental samples (four regions were sampled from two placentas each from three mice at gestational day 19) had no detectable 16S rRNA gene amplicons after PCR. Hence, more comprehensive investigations were needed.

Most recently, Younge et al. (22) used bacterial culture, fluorescent *in situ* hybridization (FISH), and 16S rRNA gene sequencing to evaluate the presence of bacterial communities in the placenta and fetal intestine of 18-30 fetuses from 2 litters in each of early, mid, and late gestations. Positive bacterial cultures were most common in mid-gestation and were not observed in late gestation. The most common bacteria cultured from the placenta and fetal intestine were *Lactobacillus, Escherichia, Enterococcus, Bacteroides*, and *Bacillus*. Mechanistic studies indicated that these cultured bacteria were not simply contaminants transferred from maternal compartments during sample processing. In the fetal intestine, bacteria were further visualized through FISH using a universal probe for the bacterial 16S rRNA gene. 16S rRNA gene profiles of the placenta and fetal intestine were similar. In early gestation, the profiles of these tissues were characterized by “*Candidatus* Arthromitus,” S24-7, *Lactobacillus*, and *Desulfovibrio*, while in mid and late gestation they were dominated by *Kurthia* and *Escherichia*. Sourcetracker analyses suggested that most of the bacterial signals from the fetal intestine in early gestation were attributed to background technical controls or to unknown sources. However, in mid and late gestation, the bacterial signals in the fetal intestine were indicated to potentially have come from the placenta or amniotic membrane. Therefore, the conclusion of the study was that there is fetal exposure to microbial communities from the placenta and the extraplacental membranes *in utero*.

### The findings of this study in the context of prior reports

In the current study, culture of bacteria from placental and fetal tissues was generally rare. Most of the bacterial isolates were identified as *Staphylococcus hominis*. The origin of these bacteria could be maternal sites, as *Staphylococcus* spp. were cultured from maternal sites and *Staphylococcus hominis* specifically was identified in molecular surveys of the maternal skin. Alternatively, these bacteria could potentially be contaminants from laboratory personnel, given that two of the five bacterial isolates recovered from negative control plates in this study were also *Staphylococcus hominis*. The other bacteria (*Bacillus, Corynebacterium, Paenibacillus*, and *Propionibacterium*) cultured from placental and fetal tissues were rarely, if ever, cultured from maternal samples or identified in the molecular surveys of maternal samples. Given that the only possible source of placental and fetal microbiota is microorganisms in the maternal compartments, the latter finding suggests that these bacteria were likely contaminants. Furthermore, there was no consistent recovery in culture of specific microorganisms (aside from *Staphylococcus hominis*) from multiple placental and fetal tissues from the same fetus or in the same tissue types among fetuses from different litters. Notably, the taxonomic identities of bacteria cultured in the current study generally differed, with the exception of *Staphylococcus* and *Bacillus*, from those initially reported by Younge et al. in placental and fetal tissues. Therefore, across current murine studies, culture has not provided consistent evidence for a placental or fetal microbiota.

In the current study, qPCR revealed that the bacterial loads of the placenta, fetal lung, liver, brain, and intestine did not exceed those of background technical controls, whereas samples from maternal sites, excluding the peritoneum, did exceed those of controls. In addition, there was no variation in bacterial load among placental and fetal tissues. These results are in contrast to those of Martinez et al. (40) in which the bacterial loads of fetal intestine exceeded those of the placenta. To our knowledge, no other studies have directly compared the bacterial loads of the placenta and fetal intestine in mammals. However, the qPCR results in our study are in agreement with prior qPCR investigations of human placental tissues – the bacterial loads of placentas are indistinguishable from those of background technical controls (7, 14, 21). Hence, there remains disagreement among studies with respect to the extent of bacterial biomass in placental and fetal tissues.

Herein, the murine placenta and fetal tissues did not yield substantive 16S rRNA gene sequence libraries, while the maternal sites other than the uterus, heart, and liver consistently did so. These results are consistent with those of Kuperman et al. (18), in which 30 cycles of PCR did not yield discernible amplicons from murine placental tissue. Notably, in our study, triple library preparations were performed and pooled for each sample, and still minimal amplicons were generated after 30 cycles of PCR. Martinez et al. (40) also used 30 cycles of PCR in their sequence library preparations and included samples in their analyses if they yielded at least 200 quality-filtered sequences, reporting a distinct bacterial DNA signal in the placenta and fetal intestine. In this study, we only included samples in analyses if they yielded at least 250 quality-filtered sequences with a Good’s coverage value of at least 95%. If we had used the criterion of 200 sequences, independent of any consideration of Good’s coverage, only one additional fetal sample would have been included in analyses. Younge et al. (22) generated substantive sequence libraries for placental and fetal intestine samples; however, their library preparation protocol was based on that of the Earth Microbiome Project (i.e. 35 cycles of PCR). The discrepancies among murine studies may therefore be due to underlying differences in the sequence library protocols used. Nevertheless, as with culture and qPCR approaches, we did not find consistent evidence of a bacterial signal in placental and fetal tissues using DNA sequencing.

Notably, in this study, there was only one case in which a bacterial isolate (i.e. *Bacillus circulans*) from a placental or fetal sample (i.e. fetal brain) had a bacterial load exceeding that of all background technical controls, and in which the bacterium was also identified in molecular surveys of at least one corresponding maternal sample (i.e. maternal skin). Therefore, in this one case, there may have been hematogenous transfer from a distant maternal site to the fetus. However, overall, there was not consistent evidence of resident bacterial communities in the murine placenta or the fetus.

### Strengths of this study

First, this study included multiple modes of microbiological inquiry, including bacterial culture, 16S rRNA gene qPCR, and 16S rRNA gene sequencing, to determine if the placental and fetal tissues of mice harbored bacterial communities. Second, this study included the analysis of many maternal, placental, and fetal body sites, including valuable low microbial biomass maternal sites such as the lung, cervix, and skin (i.e. positive controls), and maternal sites presumed to be sterile such as the liver and heart (i.e. negative controls). Third, thorough controls for potential background contamination were incorporated into bacterial culture, qPCR and DNA sequence-based analyses.

### Limitations of this study

First, this study did not include fluorescent *in situ* hybridization (FISH) to visualize potential bacterial communities in the placental and fetal tissues of mice, since protocols for low bacterial biomass FISH have not yet passed internal validation. Second, this study did not include tissue samples spiked with known numbers of bacterial cells, which could have provided information on the limits of microbial detection in the investigative approaches we used. Third, this study focused exclusively on evaluating the existence of bacterial communities in murine placental and fetal tissues; eukaryotic microbes and viruses were not considered in this study.

### Conclusion

Using bacterial culture, 16S rRNA gene qPCR, and 16S rRNA gene sequencing, there was not consistent and reproducible evidence of bacterial communities inhabiting the placenta or fetal tissues of mice, providing further evidence against the *in utero* colonization hypothesis. In addition, these findings emphasize the importance of including appropriate background technical controls, as well as positive and negative tissue controls, in all microbiological approaches from culture to sequencing when reevaluating paradigms of sterility.

